# Contextual Tumor Suppressor Function of T Cell Death-Associated Gene 8 (TDAG8) in Hematological Malignancies

**DOI:** 10.1101/144725

**Authors:** Calvin R. Justus, Edward J. Sanderlin, Lixue Dong, Alice Sun, Jen-Tsan Chi, Kvin Lertpiriyapong, Li V. Yang

**Author notes:** Address correspondence to: Li V. Yang, Ph.D., Division of Hematology/Oncology, Department of Internal Medicine, Brody School of Medicine, East Carolina University, 600 Moye Blvd., Greenville, North Carolina 27834, USA.

## Abstract

**Key points:** 1. TDAG8 (GPR65) gene expression is down-regulated in hematological malignancies.

2. Restoration of TDAG8 gene expression in blood cancer cells suppresses tumor growth and metastasis.

**Abstract:** Extracellular acidosis is a condition found within the tumor microenvironment due to inadequate blood perfusion, hypoxia, and altered tumor cell metabolism. Acidosis has pleiotropic effects on malignant progression; therefore it is essential to understand how acidosis exerts its diverse effects. In this study a bioinformatics analysis revealed the expression of the proton sensing G-protein-coupled receptor TDAG8 is significantly reduced in human blood cancers in comparison to normal blood cells. To understand how TDAG8 functions in hematological malignancies, TDAG8 expression was restored in U937 acute myeloid leukemia cells and other blood cancer cells. It was discovered that severe acidosis, pH 6.4, inhibited U937 cell proliferation while mild acidosis, pH 6.9, stimulated cell proliferation. However, restoring TDAG8 gene expression modulated the U937 cell response to mild extracellular acidosis and physiological pH by reducing cell proliferation. Tumor xenograft experiments further revealed that restoring TDAG8 expression in U937 and Ramos cancer cells reduced tumor growth. It was also shown U937 cells with restored TDAG8 expression attached less to Matrigel, migrated slower toward a chemoattractant, and metastasized less in severe combined immunodeficient mice. These effects correlated with a reduction in c-myc oncogene expression. The mechanistic investigation indicated that Gα13/RhoA signaling arbitrated the TDAG8-mediated c-myc oncogene repression in response to acidosis. Overall, this study provides compelling data to support the concept that TDAG8 functions as a contextual tumor suppressor in hematological malignancies and sensitizes blood cancer cells to acidotic stress.

## Introduction

In the early twentieth century Otto Warburg recognized a metabolic phenomenon that occurred in cancer cells, currently known as the Warburg Effect^1–4^. Warburg discovered cancer cells favor glycolysis rather than oxidative phosphorylation for energy production, even in the presence of oxygen^1–4^. It was originally hypothesized that irreversible injury of mitochondrial respiration is the cause of cancer cell formation^1^. However, this hypothesis was to some extent discredited as most cancers retain their ability to exploit mitochondrial respiration, although to a lesser degree than normal cells^5–7^. As a result of unrestricted glycolysis the tumor microenvironment is spatially acidic^8–10^. Extracellular acidosis has pleiotropic effects on tumor growth and cancer progression^10–14^. Tumor acidosis can stimulate cell death, reduce cell proliferation, and induce chromosomal instability of normal somatic cells and cancer cells^13–16^. In addition, tumor cells that become resistant to extracellular acidosis have been reported more malignant and invasive^17,18^. Therefore, extracellular tumor acidosis augments cancer progression in a Darwinian manner worsening disease prognosis. As such, it is imperative to understand how tumor cells sense and respond to acidic surroundings for adequate comprehension of cancer development.

T cell death-associated gene 8 (TDAG8, also known as GPR65) is a member of the proton sensing G-protein coupled receptor family, which also includes GPR4, GPR68 (OGR1), and GPR132 (G2A). The family of G-protein coupled receptors is activated by extracellular acidosis, which illuminates a receptor signaling connection to the acidic conditions found in tumor microenvironment^19^. The human TDAG8 gene has been mapped to chromosome 14q31-32.1, a location that abnormalities associated with T cell lymphoma and leukemia are found and is primarily expressed in immune cells and leukocyte-rich tissues such as circulating peripheral leukocytes, spleen, thymus, and tonsils^20,21^. TDAG8 has been reported to have both pro- and anti-oncogenic effects in blood cancers, which indicate TDAG8 effects are cell type and context dependent^22–24^. Therefore, it is imperative to understand the contextual effects of TDAG8 in hematological malignancies. Blood cancer cells are generally glycolytic and produce lactic acid that can acidify the microenvironment. Some types of hematological malignancies, such as lymphomas, have the ability to form solid tumors in which the tumor microenvironment is acidic. Other types of hematological malignancies, such as leukemia and multiple myeloma, form in and metastasize to bone marrow that has hypoxic and possibly acidic niches^25,26^. In rare cases, systemic lactic acidosis occurs in patients with hematological malignancies and is associated with poor prognosis^27^.

In this study the effects of acidosis and TDAG8 gene expression was investigated in blood cancers. TDAG8 gene expression was examined in hematological malignancies revealing a significant reduction in comparison to normal immune cells and leukocyte-rich tissues. Functional studies demonstrated that restoration of TDAG8 gene expression suppressed the growth, migration and metastasis of blood cancer cells and sensitized them to extracellular acidosis.

## Materials and Methods

### Bioinformatics

TDAG8 gene expression was investigated in blood cancers using the Oncomine database. A differential analysis of TDAG8 gene expression was performed between leukemia, lymphoma, and myeloma datasets and normal immune cells and leukocyte-rich tissue. Additional bioinformatics analyses were performed on datasets from the National Center for Biotechnology Information (NCBI) Gene Expression Omnibus (GEO). The raw data was downloaded using expression console software HG_u133_Plus_2 and HG_u133a_2 as libraries. The analysis was run and the RMA normalization method was used to generate the values graphed. The output file was then merged with the probe set downloaded from Affymetrix.

### Plasmid Constructs

The MSCV-huTDAG8-IRES-GFP construct was made previously and the empty construct, MSCV-IRES-GFP, was used as a control^23^. For the pMSCV-huTDAG8-IRES-YFP-P2A-OF-LUC construct, the open reading frame of human TDAG8 was amplified using primers designed to contain the EcoRI and XhoI restriction enzyme sites: 5′-ATAAGAAT<u>GAA</u> <u>TTC</u>ACCATGAACAGCACATGTATTGAAGAA-3′ and 5′-ATAAGAATGAATTC<u>CTCGAG</u> CTACTCAAGGACCTCTAATTCCAT-3′. The pMSCV-IRES-YFP-P2A-OF-LUC plasmid was then digested with EcoRI and XhoI and the huTDAG8 open reading frame was cloned into it generating the pMSCV-huTDAG8-IRES-YFP-P2A-OF-LUC construct. The resultant construct was verified by DNA sequencing.

### Cell Lines and Culture

All cell lines were cultured in Roswell Park Memorial Institute medium (RPMI) supplemented with 10% fetal bovine serum (FBS) in a tissue culture incubator set at 37°C and 5.0% CO_2_. RPMI/FBS medium was buffered using 7.5 mM HEPES, 7.5 mM EPPS and 7.5 mM MES, known collectively as RPMI/HEM, as previously described^23^. Cells used for experiments were >95% viable as assessed by the trypan blue dye exclusion method.

To generate green fluorescent protein (GFP) and yellow fluorescent protein (YFP)/luciferase (Luc) cell lines with restored TDAG8 gene expression, retroviral-mediated cell transduction was performed as previously described^28^. To generate the Gα13 signaling-deficient cell lines the p115 Rho RGS construct was subcloned into the pQCXIP vector as previously described^28,29^. The p115 Rho RGS-pQCXIP and empty pQCXIP retroviral vectors were then stably transduced into U937 cells.

### EdU Cell Proliferation Assay

The Click-iT® Plus EdU Imaging Kit was used to examine U937/Vector and U937/TDAG8 cell proliferation. Fluorescence microscopy with the EVOS®*fl* Digital Inverted Microscope was used to take images of cells incorporating Hoescht® 33342 dye and the EdU analogue, and also images with transmitted light in the same field of view at a 200× magnification. Adobe Photoshop’s counting tool was used to count the number of total cells according to Hoescht® 33342 dye and proliferating cells according to EdU positive cells.

### Cell Growth Competition Assay

The same number of non-GFP expressing cells (U937/Parental or RPMI 8226/Parental) and GFP expressing cells (Vector or TDAG8) were mixed into a well and the percentages of non-GFP and GFP cells were measured using flow cytometry over 14 days. Cell population percentages were analyzed and graphed.

### Annexin V/7AAD Staining

Cells were stained with the PE Annexin V Apoptosis Detection Kit I (BD Biosciences). Emission of single cell fluorescence was measured at 572 nm for Annexin V, 647 nm for 7AAD, and 525 nm for GFP. The results were analyzed with CellQuest software (BD Biosciences).

### Quantitative reverse transcription-polymerase chain reaction (RT-PCR)

Gene expression was determined by quantitative RT-PCR as previously described^23^, using the following TaqMan predesigned primer/probes from Invitrogen, TDAG8 (Hs00269247_s1), c-myc (Hs00153408_m1), and β-actin (Hs99999903_m1). Relative gene expression was calculated using the 2^−ΔΔCt^ method^30^.

### Western Blot

Western blot was performed as previously described^23^. Antibodies of c-myc (product #5605), phospho-paxillin (Y118) (#2541), phospho-CREB (S133) (#9198), CREB (#9197), caspase-3 (#9665), caspase-9 (#9502), cleaved PARP XP (#5625), and β-actin (#4970) were purchased from Cell Signaling Technology.

### Histology

Immunohistochemistry for antibodies against c-myc (Abcam, #ab32072), cleaved PARP (Cell Signaling Technology, #5625), Ki67 (Abcam, #ab15580), and human nucleoli (Abcam, #ab190710) was performed on paraffin-embedded 5μm sections. Antigen retrieval was performed by boiling slides in TRIS-EDTA + 0.1% Tween-20 pH 9.0 antigen retrieval buffer for 10-18 minutes. SuperPicture^™^ 3rd Gen IHC Detection Kit (Invitrogen) was used to complete immunohistochemistry.

### Transwell Cell Migration Assays

Two hundred μl of cells were seeded into the transwell insert at 5× 10^6^ cells/ml in migration buffer consisting of RPMI media buffered to pH 7.4 and pH 6.4 without FBS and supplemented with 0.1% BSA. Chemoattractant was then added into the bottom well. The plates were then incubated at 37° C and 5% CO_2_ for 5 hours. After 5 hours the number of migrated cells was quantified using flow cytometry.

### Cell Attachment Assays

Matrigel solution was added into each well to form a thin layer of gel. U937 cells were plated onto Matrigel and incubated at 37°C and 5% CO_2_ for 1 hour in culture media buffered to pH 7.4 or 6.4. After 1 hour the media was aspirated and the wells were washed 4 times with RPMI to remove non-adherent cells. For the endothelial cell attachment assay Human Umbilical Vein Endothelial cells (HUVEC) were seeded on 0.1% gelatin-coated plates to create a monolayer. Next, U937 cells were plated in each well in culture media buffered to pH 7.4 or 6.4. Plates were incubated at 37°C and 5% CO_2_ for 1 hour. After 1 hour the media was aspirated and the wells were washed 4 times with RPMI. For all attachment assays several images were taken in various areas of the well to give an adequate representation. Cell attachment was quantified by counting every cell in each field of view.

### Primary Tumor and Metastasis Xenograft

NOD.CB17-Prkdc<SCID>/J mice were purchased from Jackson Laboratories and bred at East Carolina University animal facilities for research purposes. All experimental procedures were performed according to institutional regulations. In each experiment, 5× 10^6^ U937 or Ramos cells were injected into the flanks of SCID mice. Tumor growth was measured daily using a caliper. Tail vein injections were performed with 2× 10^6^ U937 cells in SCID mice. When mice reached endpoint parameters, e.g. lethargy, hunched posture, and unkempt appearance, they were euthanized for analysis. The experiment was terminated 8 months after injection when the remaining mice showed no signs of disease progression. To monitor U937-Luc tumor growth in vivo, intraperitoneal injection with D-Luciferin (Caliper Life Sciences, Catalogue # 119222) at 15 mg/ml in DPBS was performed. Subsequently, mice were imaged using the IVIS Lumina XR unit.

### Statistical Analysis

Statistical analysis was performed using the GraphPad Prism software. The unpaired *t*-test was used to test for statistical significance between 2 groups. Error bars represent ± standard error (S.E.M.) in all graphs. ns P > 0.05, * P < 0.05, ** P < 0.01, *** P < 0.001.

## Results

### TDAG8 gene expression is reduced significantly in hematologic malignancies in comparison to normal immune cells and leukocyte-rich tissue

Oncomine bioinformatics analysis identified that TDAG8 gene expression was reduced in multiple forms of blood cancer, including acute myeloid leukemia (AML) (2.2 and 6.1 fold)^31,32^, chronic lymphocytic leukemia (CLL) (3.3, 1.7, 2.3, and 2.6 fold) (Table 1)^31,33–35^, T-cell acute lymphoblastic leukemia (TCALL) (3.1 fold), B-cell childhood acute lymphoblastic leukemia (BCCALL) (2.5 fold), B-cell acute lymphoblastic leukemia (BALL) (2.4 fold), pro B-cell acute lymphoblastic leukemia (PBALL) (1.3 fold), hairy cell leukemia (HCL) (1.9 fold), and T-cell prolymphocytic leukemia (TCPLL) (3.9 fold) (Table 1)^31,33,36^. In addition, TDAG8 gene expression is reduced in several lymphomas such as diffuse large B-cell lymphoma (DLBCL) (1.4 and 1.9 fold)^33,34^, follicular lymphoma (FL) (1.4 and 2.0 fold)^33,34^, pleural effusion lymphoma (PEL) (2.7 fold)^33^, and Burkitt lymphoma (BL) (1.8 fold)^33^ (Table 1). Lastly, TDAG8 expression was found reduced in smoldering myeloma (SM) (2.6 fold) (Table 1)^37^. The search in the Oncomine^™^ Research Premium Edition also revealed various cancer cell lines that were resistant to chemotherapeutics had reduced TDAG8 gene expression in comparison to cancer cell lines that were sensitive to those particular chemotherapeutics (Table 2)^38,39^. A separate analyses of datasets from NCBI GEO revealed that TDAG8 gene expression is significantly reduced 4.5 fold in AML (GEO Series ID GSE9476), 2.9 fold in CLL (GEO Series ID GSE22529), 3.6 fold in TCPLL with inv(14)(q11q32) (GEO Series ID GSE5788), 2.7 fold in chronic B-cell lymphocytic leukemia (CBLL) (GSE26725), 3.0 fold in DLBCL (GEO series ID GSE12195), and 2.8 fold in FL (GEO series ID GSE12195) (Figure 1A-E). The same pattern was found in various blood cancer cell lines. For example, TDAG8 gene expression was reduced 45 fold in U937 cells, 8 fold in Ramos cells, undetectable in RPMI 8226 cells, 625 fold in K562 cells, and 3 fold in Jurkat cells in comparison to normal peripheral blood leukocytes (Figure 2A).

**Figure 1.**
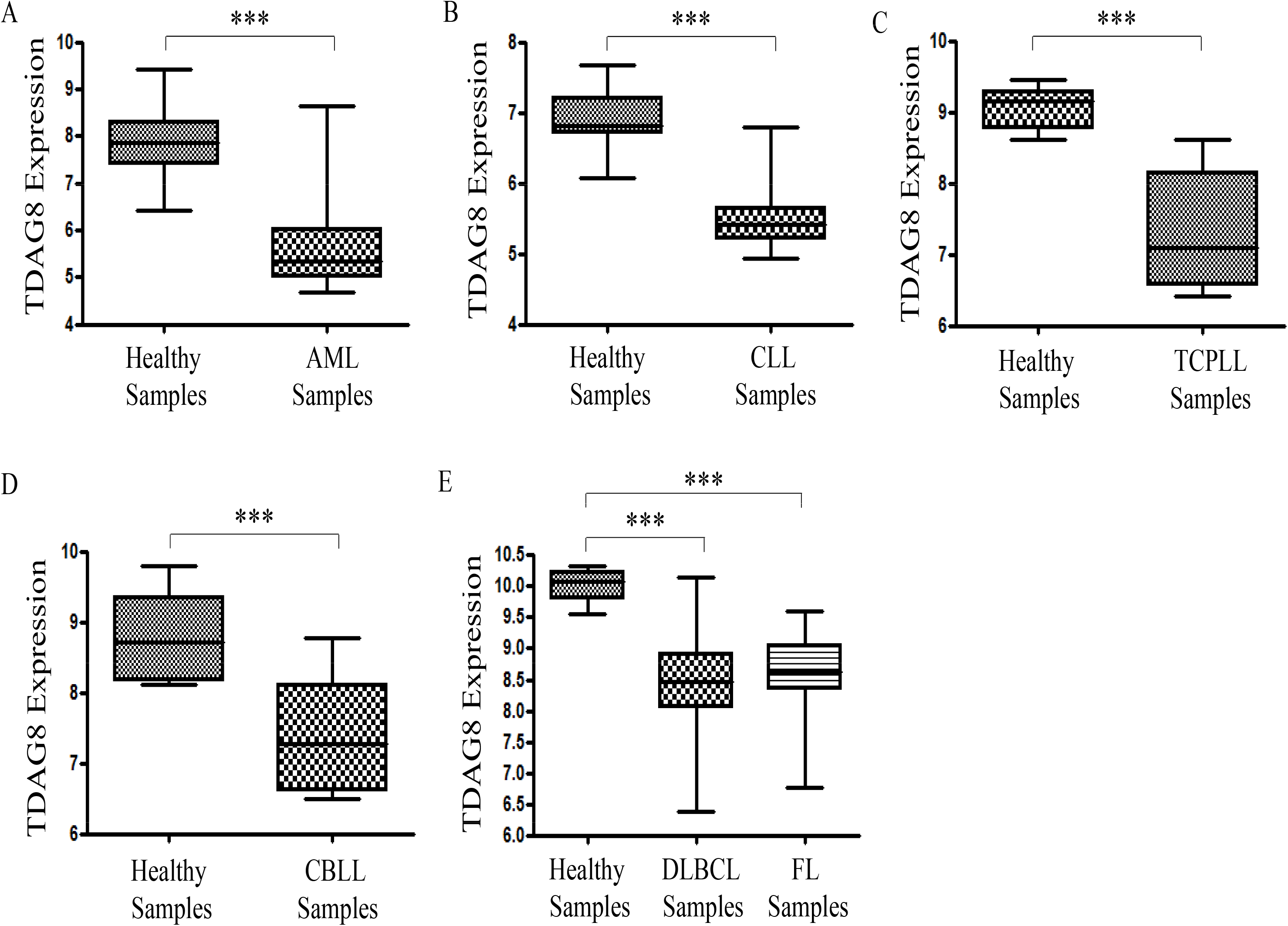
TDAG8 gene expression is reduced in patients with blood cancers in comparison to healthy donor’s immune cells or leukocyte-rich tissue. (A) The expression of TDAG8 is reduced 4.5 fold in CD34+ bone marrow and CD34+ peripheral blood cells isolated from patients with AML (N=20) in comparison to CD34+ bone marrow and CD34+ peripheral blood cells isolated from healthy donors (N=26). (B) The expression of TDAG8 is reduced 2.9 fold in CD19+/CD5+ peripheral blood mononuclear cells isolated from patients with CLL (N=11) in comparison to CD19+/CD5+ peripheral blood mononuclear cells isolated from healthy donors (N=39). (C) The expression of TDAG8 is reduced 3.6 fold CD3+ peripheral blood cells (N=8) isolated from patients with TCPLL in comparison to CD3+ peripheral blood cells isolated from healthy donors (N=6). (D) The expression of TDAG8 is reduced 2.7 fold in CD19+ peripheral blood lymphocytes isolated from patients with CBLL (N=6) in comparison to CD19+ peripheral blood lymphocytes isolated from healthy donors (N=11). (E) The expression of TDAG8 is reduced by 3.0 fold in DLBCL biopsy samples (N=73) in comparison to B-lymphocytes isolated from healthy donors (N=9). The expression of TDAG8 is also reduced by 2.8 fold in human FL biopsy samples (N=38) in comparison to B-lymphocytes isolated from healthy donors (N=9). AML-acute myeloid leukemia, CLL-chronic lymphocytic leukemia, DLBCL-diffuse large B-cell lymphoma, FL-follicular lymphoma, TCPLL-T-cell prolymphocytic leukemia with inv(14)(q11q32), CBLL-chronic B-cell lymphocytic leukemia. Y-axis is Log_2_ scale. *** P < 0.001

**Figure 2.**
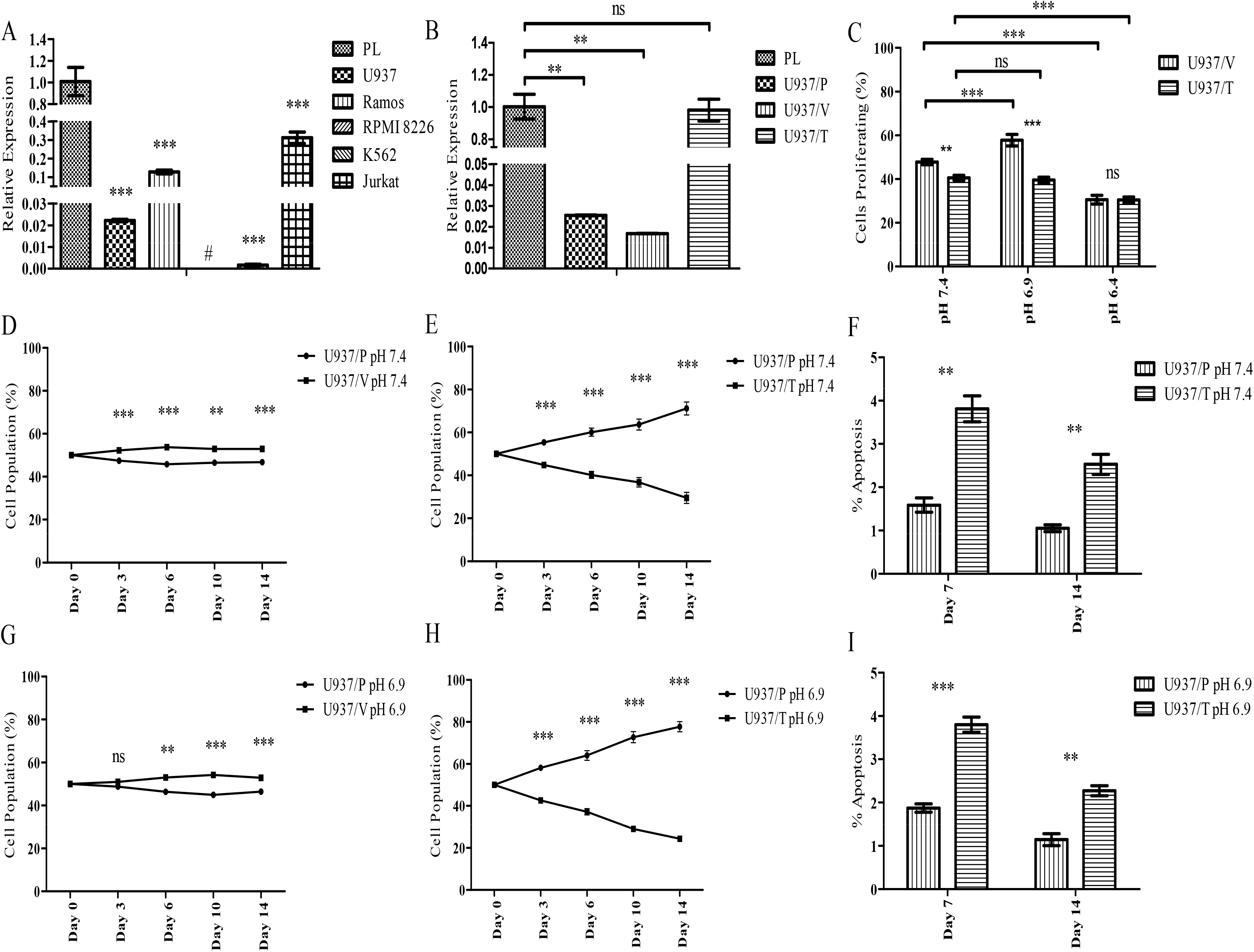
Restoration of TDAG8 gene expression in U937 cells provides a growth disadvantage at physiological pH 7.4 and mildly acidic pH 6.9. (A) Quantitative RT-PCR demonstrates a reduction of TDAG8 gene expression in U937, Ramos, RPMI 8226, K562, and Jurkat cells in comparison to peripheral leukocytes pooled from 426 healthy donors. (B) Quantitative RT-PCR demonstrates restoration of TDAG8 gene expression in U937 acute myeloid leukemia cells following transduction and cell sorting. (C) Mild acidosis (pH 6.9) increases U937/Vector cell proliferation while severe acidosis (pH 6.4) inhibits cell proliferation. Restoring TDAG8 gene expression in U937 cells inhibits cell proliferation in comparison to U937/Vector cells. (D) The expression of the empty MSCV-IRES-GFP construct in U937 cells does not substantially affect the U937/Vector cell population percentage in comparison to U937/Parental cells at pH 7.4 from day 0 to day 14. (E) The restoration of TDAG8 gene expression in U937 cells reduces cell growth in comparison to U937/Parental cells at physiological pH 7.4. (F) Restoring TDAG8 gene expression in U937 cells increases apoptosis throughout the cell competition assay at pH 7.4. (G) The expression of the empty MSCV-IRES-GFP construct in U937 cells does not substantially affect the U937/Vector cell population percentage in comparison to U937/Parental cells at pH 6.9 from day 0 to day 14. (H) The difference between the U937/TDAG8 cell population and the U937/Parental cell population throughout the competition assay at acidic pH 6.9 is greater in comparison to pH 7.4 treatment. (I) Restoring TDAG8 gene expression in U937 cells increases apoptosis throughout the cell competition assay at pH 6.9. Pooled RNA from 426 normal peripheral leukocytes was purchased from Clontech Laboratories, Inc. to be used as a control for a healthy immune cell comparison. ns P > 0.05, * P < 0.05, ** P < 0.01, *** P < 0.001

**Table 1.**
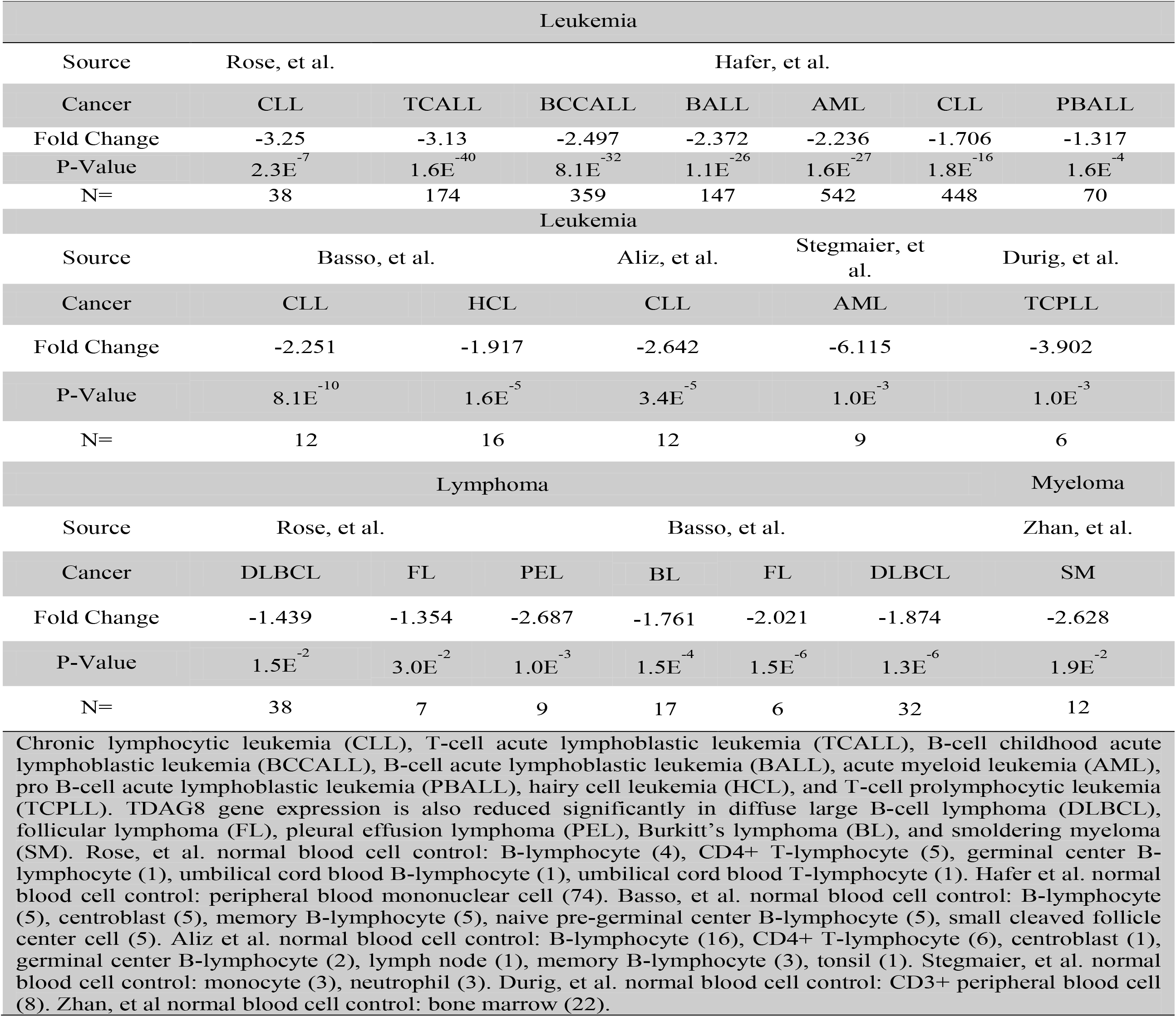
TDAG8 gene expression is reduced significantly in blood cancers

**Table 2.** Reduced TDAG8 gene expression correlates with resistance to various chemotherapeutics

| Drug | Elesclomol<br>(Leukemia) | Panobinostat<br>(Multi-Cancer) | Irinotecan<br>(Multi-Cancer) | Topotecan<br>(Multi-Cancer) | Navitoclax<br>(Leukemia) | MK2206<br>(Leukemia) |
| --- | --- | --- | --- | --- | --- | --- |
| Mechanism<br>of Action | Unknown<br>Oxidative<br>Stress | Histone<br>Deacetylase<br>Inhibitor | Topoisomerase<br>Inhibitor | Topoisomerase<br>Inhibitor | Bcl-2, Bcl-XL, and Bcl-<br>w Inhibitor | AKT<br>Inhibitor |
| Resistant<br>Cell Lines<br>(*Rn*) | 21 | 164 | 84 | 124 | 12 | 17 |
| Sensitive<br>Cell Lines<br>(*Sn*) | 4 | 86 | 71 | 154 | 6 | 3 |
| TDAG8<br>Gene<br>Expression<br>of *Sn* vs *Rn* | +2.3 | +2.7 | +3.2 | +2.2 | +2.4 | +3.8 |
| P Value | 4.8E<sup>-2</sup> | 2.6E<sup>-7</sup> | 4.2E<sup>-8</sup> | 2.9E<sup>-10</sup> | 3.4E<sup>-2</sup> | 7.8E<sup>-4</sup> |

Cancer cells of various origins categorized as resistant to specific chemotherapeutics demonstrate a significant reduction in TDAG8 gene expression. The expression of TDAG8 in 4 cell lines sensitive to elesclomol is 2.3 fold higher than in 21 cell lines resistant. The expression of TDAG8 in 86 cell lines categorized as sensitive to panobinostat is 2.7 fold higher than in 164 cell lines categorized as resistant. The expression of TDAG8 in 71 cell lines categorized as sensitive to irinotecan is 3.2 fold higher than in 84 cell lines categorized as resistant. The expression of TDAG8 in 154 cell lines categorized as sensitive to topotecan is 2.2 fold higher than in 124 cell lines categorized as resistant. The expression of TDAG8 in 6 cell lines categorized as sensitive to navitoclax is 2.4 fold higher than in 12 cell lines categorized as resistant. The expression of TDAG8 in 3 cell lines categorized as sensitive to MK2206 is 3.8 fold higher than in 17 cell lines categorized as resistant.

### TDAG8 gene expression provides a selective disadvantage to blood cancer cell growth

As TDAG8 gene expression is commonly downregulated in hematological malignancies (Table 1–2 and Figure 1), we investigated its function by restoring TDAG8 expression in blood cancer cell lines such as U937, Ramos and RPMI 8226 with low TDAG8 expression. The U937 cell line was originally established from a patient with histiocytic lymphoma and later found to possess characteristics of acute myeloid leukemia (AML) cells^40^. U937 cells were used as a model system for this study because the cells have features of both lymphoma and leukemia. Proliferation of U937 cells transduced with a construct containing the human TDAG8 gene (U937/TDAG8) was compared to U937 cells transduced with an empty construct (U937/Vector). The expression level of TDAG8 mRNA in U937/TDAG8 cells was restored to the similar level as that in normal leukocytes (Figure 2B). Due to the lack of a reliable TDAG8 antibody, we were unable to detect TDAG8 protein expression. Instead, we examined the TDAG8 downstream signaling activities and found the phosphorylation of CREB was higher in U937/TDAG8 cells than that in U937/Vector cells (Supplemental Figure 1A), suggesting that restoration of TDAG8 expression augments its downstream G protein signaling. Similarly, CREB phosphorylation was increased in Ramos/TDAG8 cells with the restoration of TDAG8 expression (Supplemental Figure 1B-C).

In comparison to U937/Vector cells, U937/TDAG8 cells proliferated 7% less (47.8% vs. 40.5% proliferating cells) at physiological pH 7.4 (Figure 2B-C). When U937/Vector cells were treated with pH 6.9, there was a 10% increase in cell proliferation in comparison to physiological pH 7.4. However, restoring TDAG8 gene expression sensitized U937 cells to media buffered to pH 6.9 and suppressed cell proliferation at pH 6.9 (Figure 2C). Severe acidosis, pH 6.4, significantly reduced cell proliferation in both U937/Vector and U937/TDAG8 cells (Figure 2C).

Next, cell growth competition assays were performed to demonstrate negative selection of U937 cells expressing the TDAG8 gene. In the cell growth competition assay there was a slight change in the U937/Parental-U937/Vector cell populations (50 ±5%) after 14 days (Figure 2D). However, the U937/TDAG8 cell population was only 30% of the total population of cells in comparison to 70% of the U937/Parental cells at pH 7.4 (Figure 2E). Quantitative RT-PCR also demonstrated that restoration of TDAG8 expression in U937/TDAG8 cells reduced the expression of c-myc oncogene on day 0 and day 14 of the competition assay in comparison to the U937/Vector cells (Supplemental Figure 2A-B). The growth competition assay was also performed in medium buffered to mildly acidic pH 6.9. The U937/TDAG8 cell population was significantly reduced to 23% of the total population of cells at pH 6.9 (Figure 2H). The U937/Vector and U937/Parental cells showed slight changes (50 ±5%) relative to each other at pH 6.9 (Figure 2G). Results were similar when the competition assay was performed with the RPMI 8226/Vector and RPMI 8226/TDAG8 myeloma cells (Supplemental Figure 3).

The GFP mean fluorescence value remained stable from day 0 to day 14 with little variation in the U937/Vector cells when co-cultured with the U937/Parental cells (Supplemental Figure 2C). However, U937/TDAG8 cell GFP mean fluorescence value as an indicator of TDAG8 transgene expression was considerably reduced (Supplemental Figure 2C). When comparing day 0 with day 14 at pH 6.9 there was a 65% reduction in GFP mean fluorescence value of U937/TDAG8 cells (Supplemental Figure 2D).

Annexin V PE/7AAD staining was performed at day 7 and day 14 of the competition assay to examine U937 cell apoptosis. It was determined there was a significant increase of apoptosis in U937/TDAG8 cells in comparison to U937/Parental cells at pH 7.4 and at pH 6.9 (Figure 2F & I). Increased apoptosis of U937/TDAG8 cells also correlated with an increase in caspase 3, caspase 9, and PARP cleavage (Supplemental Figure 4).

### Restoration of TDAG8 gene expression reduces primary tumor growth in severe combined immunodeficient (SCID) mice

U937 cells were originally isolated from a patient with histiocytic lymphoma and can form solid tumors, which are known to be spatially acidic. To determine the effects of restoring TDAG8 gene expression on U937 primary tumor growth, subcutaneous xenograft experiments were performed in SCID mice. A significant inhibition of U937/TDAG8 tumor growth was observed throughout the experiments (Figure 3A). In addition, overall tumor mass was reduced by ~50% in U937/TDAG8 tumors (Figure 3B). Similar trends were found following Ramos/Vector and Ramos/TDAG8 lymphoma cell injections into SCID mice (Supplemental Figure 5). In vivo luminescence imaging was also used to visualize tumor growth over 2 weeks. The pattern previously observed in U937/TDAG8 cells, i.e. reduced tumor growth, was consistent with the U937/TDAG8-Luc cells expressing TDAG8 and the luciferase marker gene (Figure 3C).

**Figure 3.**
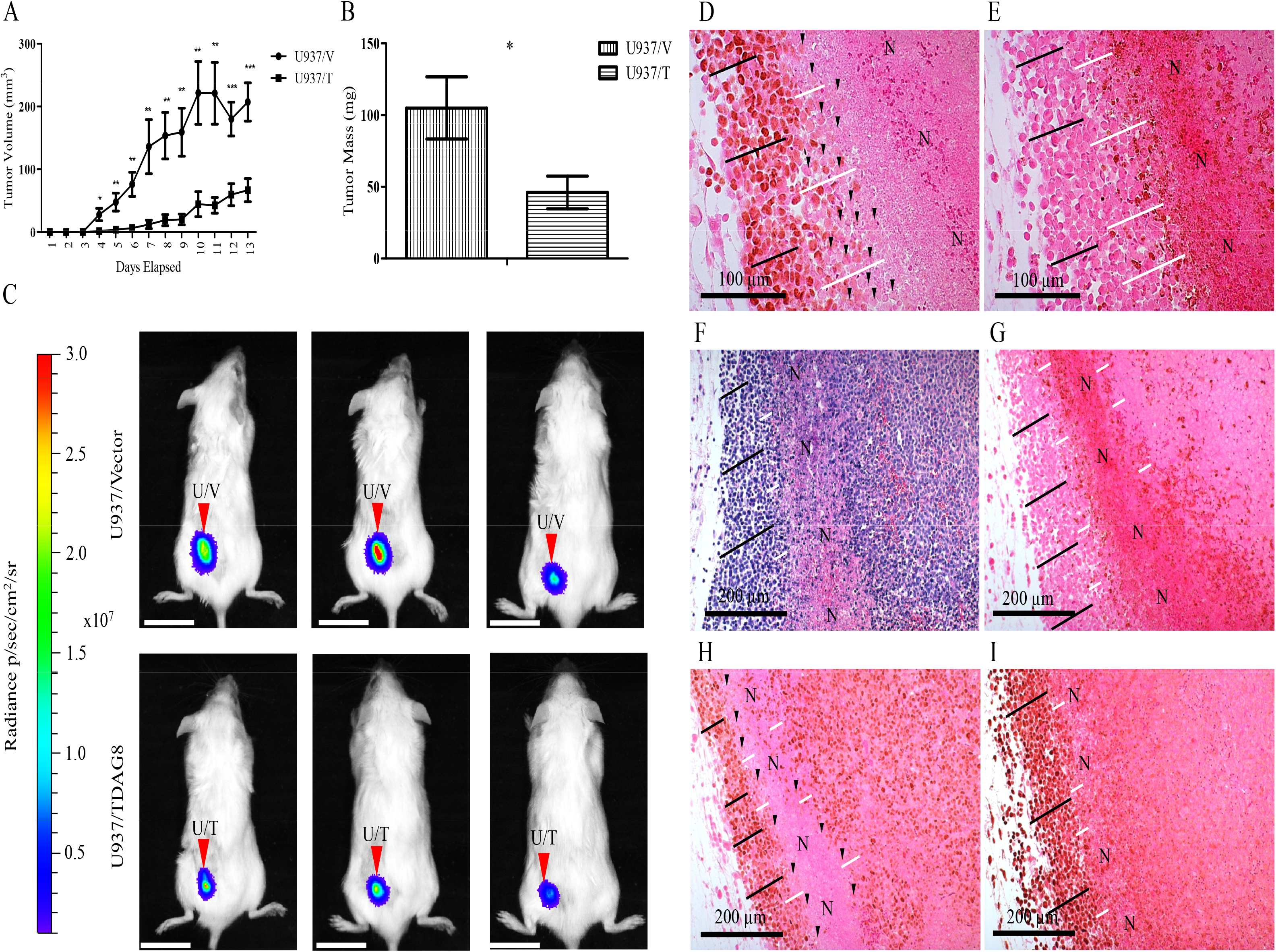
Restoring TDAG8 gene expression in U937 cells reduces primary tumor growth in SCID mice. (A) U937/TDAG8 tumor growth was significantly reduced in comparison to U937/Vector tumors from day 4 to necropsy (N=14). (B) U937/TDAG8 tumor mass was reduced significantly in comparison to U937/Vector tumors following necropsy. (C) In vivo imaging of 3 mice injected with U937/Vector-Luc or U937/TDAG8-Luc cells 6 days after injection. Scale Bar = 2 cm. (D-E) Immunohistochemistry of c-myc (D) and cleaved PARP (E) in U937/Vector tumors demonstrating regions nearest necrotic areas display reduced c-myc oncogene expression while still live. (F-I) Hematoxylin and eosin staining (F) and immunohistochemistry of U937/Vector tumors demonstrating invasive peripheral regions of U937 tumors display increased proliferation by c-myc (H) and Ki67 (I) while demonstrating less apoptosis, cleaved PARP (G). N-Necrotic. White lines indicate areas adjacent to necrotic zones. Black lines indicate tumor cells that are invasive correlating with higher c-myc and Ki67 expression. Black arrowheads indicate single tumor cells that demonstrate reduced or no c-myc expression. * P < 0.05, ** P < 0.01, *** P < 0.001

### Tumor acidosis congruent to necrotic regions inhibits c-myc expression

Immunohistochemistry (IHC) of c-myc expression revealed that sections nearest necrotic regions of tumors, known to be acidic^17^, had reduced expression of c-myc (Figure 3D-I). The areas that were investigated for low c-myc expression were negative for cleaved PARP indicating that intact cells nearest necrotic regions were viable (Figure 3E & 3G). In addition, it was also revealed that invasive cancer cells in the tumor peripheral regions had increased expression of c-myc and Ki-67 and were negative for cleaved PARP (Figure 3G-I).

### Acidosis stimulates Ga13/RhoA signaling and reduces c-myc expression in U937 cells partially through TDAG8

IHC analysis of c-myc protein expression revealed that U937/TDAG8 tumor cells expressed lower levels of c-myc in comparison to U937/Vector control (Figure 4B). It was also qualitatively determined that c-myc expression was undetectable in numerous U937/TDAG8 cells (Figure 4B). In comparison, the majority of U937/Vector cells expressed c-myc at a high level (Figure 4A). It was also determined that restoring TDAG8 gene expression reduces c-myc protein expression significantly in comparison to the U937/Vector control cells at physiological pH 7.4 (Figure 4C-D). In addition, activation of TDAG8 by extracellular acidification, representing similar conditions as the tumor microenvironment, further reduces c-myc expression in comparison to pH 7.4 treatment and to the U937/Vector controls (Figure 4C-D). These observations are consistent with previous studies showing that TDAG8 has some constitutive activities at the physiological pH and is fully activated under acidic pH^41^.

**Figure 4.**
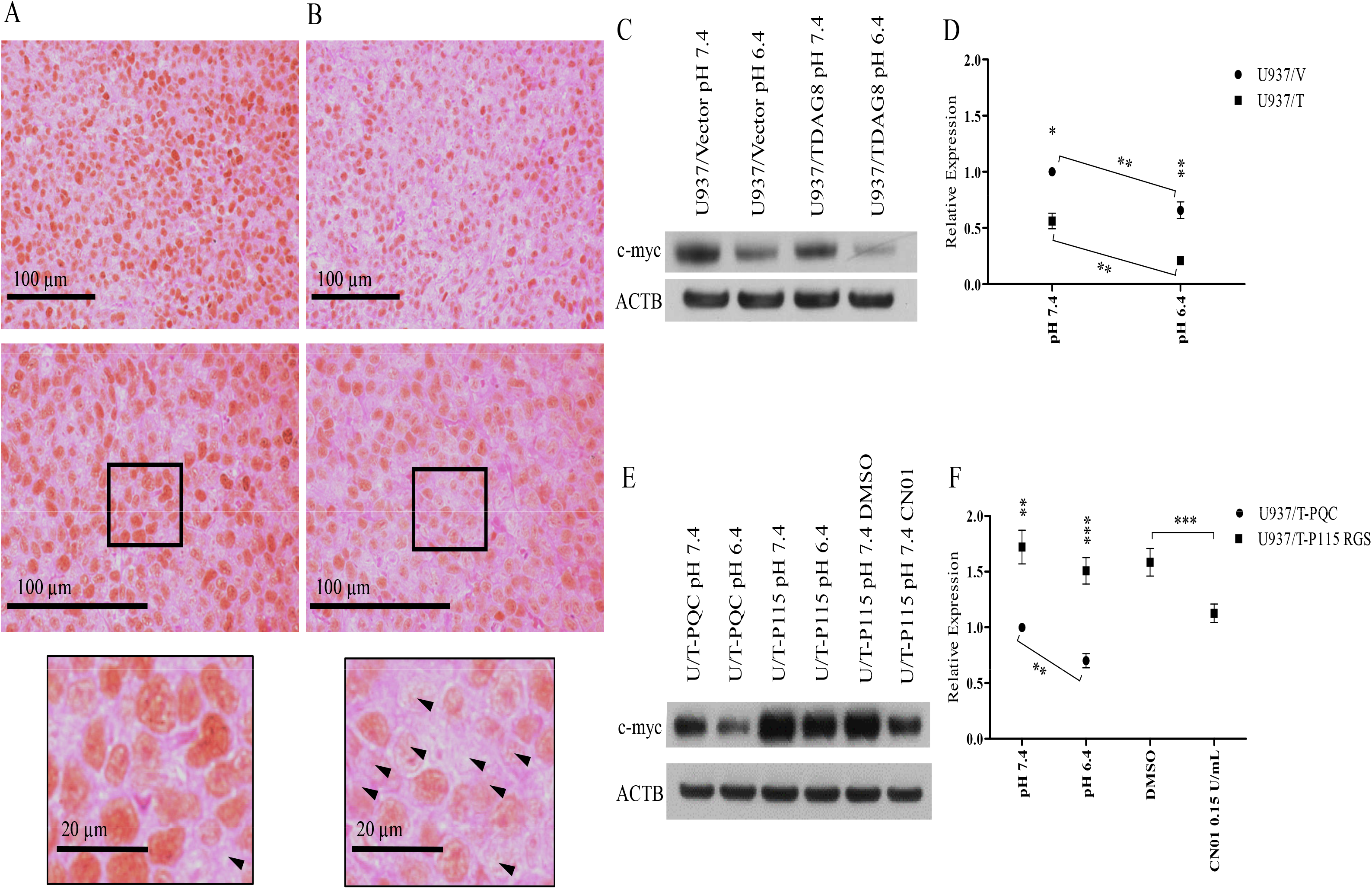
Restoration of TDAG8 gene expression reduces c-myc oncogene expression through Gα13/RhoA signaling. (A) IHC demonstrates the majority of cells within the U937/Vector tumors away from necrotic or heterotypic areas are positive for c-myc expression as indicated by DAB chromogen (N = 5). (B) There were less cells within the U937/TDAG8 tumors away from necrotic or heterotypic areas positive for c-myc expression as indicated by DAB chromogen. Arrowheads indicate cells not expressing c-myc. Observed in at least 3, 5μm sections of 5 different tumors for each experimental group. (C) Restoration of TDAG8 in U937 cells reduces c-myc expression in response to treatment with media buffered to physiological pH 7.4 and acidic pH 6.4. (D) Quantification of c-myc Western blot data displaying the significant inhibition of c-myc mediated by acidosis and TDAG8 gene expression. (E-F) Gα13/RhoA signaling is responsible for a TDAG8 mediated inhibition of c-myc expression. * P < 0.05, ** P < 0.01, *** P < 0.001

Following inhibition of Gα13 signaling by the p115 RGS construct^28,29^, Western blot analysis determined that c-myc expression was restored in the U937/TDAG8 cells after treatment with media buffered to pH 7.4 and pH 6.4 (Figure 4E-F). These results reveal that Gα13 signaling is central for TDAG8-mediated c-myc inhibition at physiological pH 7.4 and acidic pH 6.4. The activation of RhoA with CN01, downstream from the p115-RGS Gα13 inhibition, reduced the expression of c-myc, indicating that Gα13/RhoA signaling in U937 cells inhibits c-myc expression (Figure 4E-F). Similar results were found for caspase 3, caspase 9, and PARP cleavage following inhibition of Gα13 signaling and activation of RhoA (Supplemental Figure 6). Therefore, the data suggests TDAG8 exerts its growth inhibitory effects through activation of Gα13/RhoA signaling in U937 cells.

### Restoration of TDAG8 gene expression in U937 cells reduces cell attachment and migration correlating with altered focal adhesion dynamics

Restoration of TDAG8 gene expression reduced the ability for U937 cells to attach to Matrigel at physiological pH 7.4 by more than 50% (Figure 5A). Activation of TDAG8 by extracellular acidosis reduced U937 cell attachment even further demonstrating the antimetastatic capabilities of TDAG8 (Figure 5A). U937 cell attachment to a HUVEC monolayer was also investigated revealing a reduction in U937/TDAG8 cell attachment to endothelial cells at physiological pH 7.4 and acidic pH 6.4 (Supplemental Figure 7A). However, overall cell attachment to a HUVEC monolayer was increased at acidic pH 6.4 likely due to the effects of acidosis on endothelial cell-mediated attachment^42^. It was also determined that U937/TDAG8-Luc cells exhibited less cell migration towards the chemoattractant SDF-1α in comparison to U937/Vector-Luc cells (Figure 5B). In addition, treatment with pH 6.4 further reduced the migration of U937/TDAG8-Luc cells in comparison to the U937/Vector-Luc cells (Figure 5B). This was also observed in the U937 cells that were transduced with the GFP construct (Supplemental Figure 7B).

**Figure 5.**
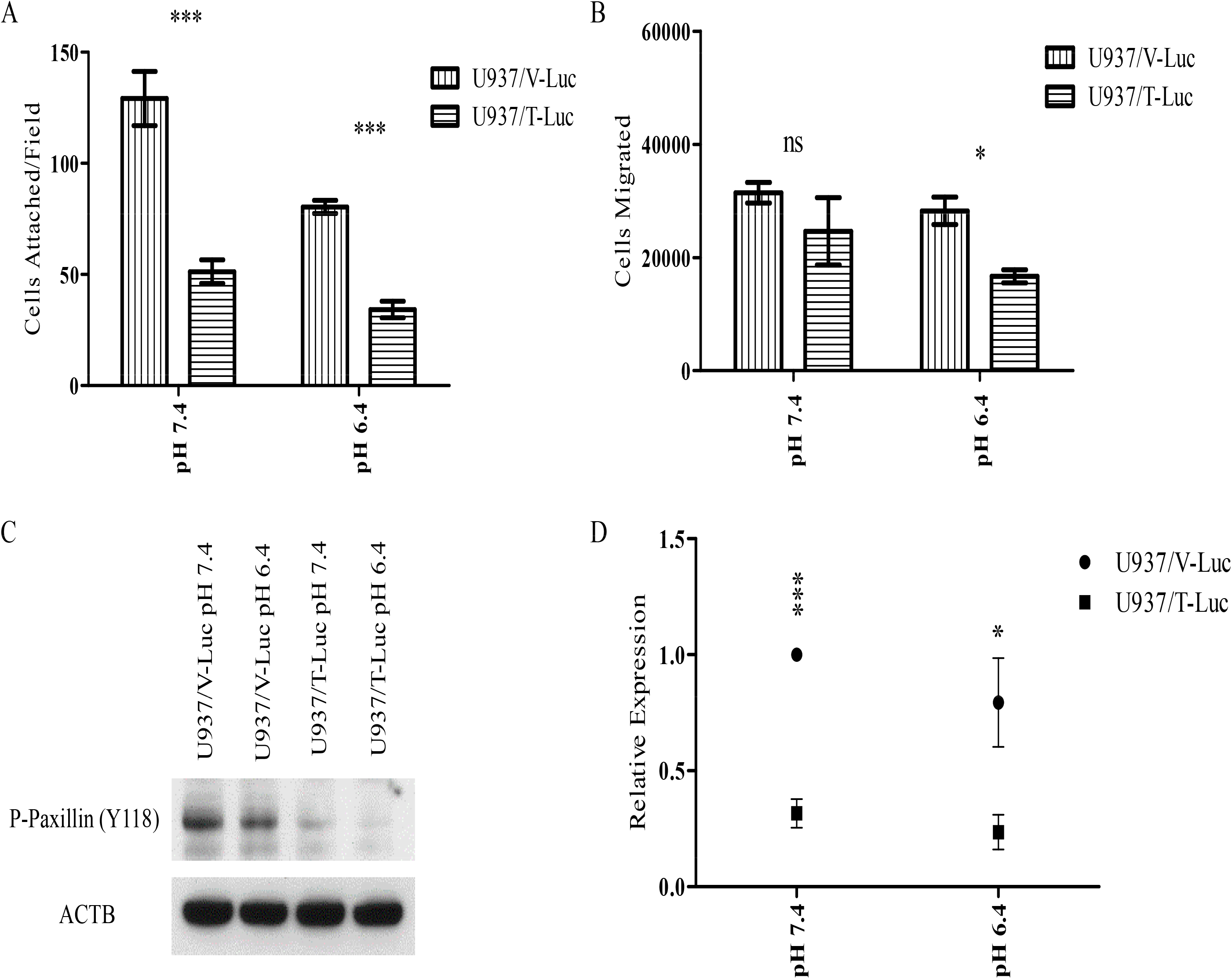
TDAG8 gene expression reduces U937 cell attachment and migration. (A) Restoration of TDAG8 gene expression inhibits U937 cell attachment to Matrigel. (B) U937 cell migration towards SDF1-α is inhibited by TDAG8 gene expression at pH 7.4 and pH 6.4. (C) Acidosis and restoration of TDAG8 gene expression reduces phosphorylation of paxillin at Y118 in U937 cells. (D) Quantification of paxillin phosphorylation at Y118 reveals a significant reduction in U937/TDAG8-luc cells at pH 7.4 and 6.4. ns P > 0.05, * P < 0.05, ** P < 0.01, *** P < 0.001

Increased activity of focal adhesion proteins such as focal adhesion kinase (FAK) and paxillin have been correlated with poor patient survival and increased malignancy of blood and other cancers^43–48^. We found that restoring TDAG8 gene expression in U937 cells significantly inhibited the phosphorylation of paxillin Y118 at pH 7.4 and pH 6.4 (Figure 5C-D), suggesting a reduced activity of the FAK/paxillin pathway^49,50^. In addition, when treated with media buffered to pH 6.4 the phosphorylation of paxillin Y118 was reduced in comparison to the pH 7.4 treatment groups (Figure 5C-D).

### Restoration of TDAG8 gene expression in U937 cells reduces metastasis in SCID mice following tail vein injections

Overall, the in vivo imaging of SCID mice was not sensitive enough to visualize micrometastasis of tumor cells (Figure 6A-E). However, larger nodules and tumors were easily visualized (Figure 6A). Metastasis of U937/Vector cells to the lymph node, bone of the limb, and other organs in SCID mice was detected by IVIS luminescence imaging (Figure 6A & Supplemental Figure 7C). To further quantify metastasis, the mouse lungs were analyzed using an anti-human nucleoli antibody specific for human cells (Figure 6B-E). The results indicate the severity of U937 cell metastasis to the lung of SCID mice was significantly reduced by restoring TDAG8 expression (Figure 6F).

**Figure 6.**
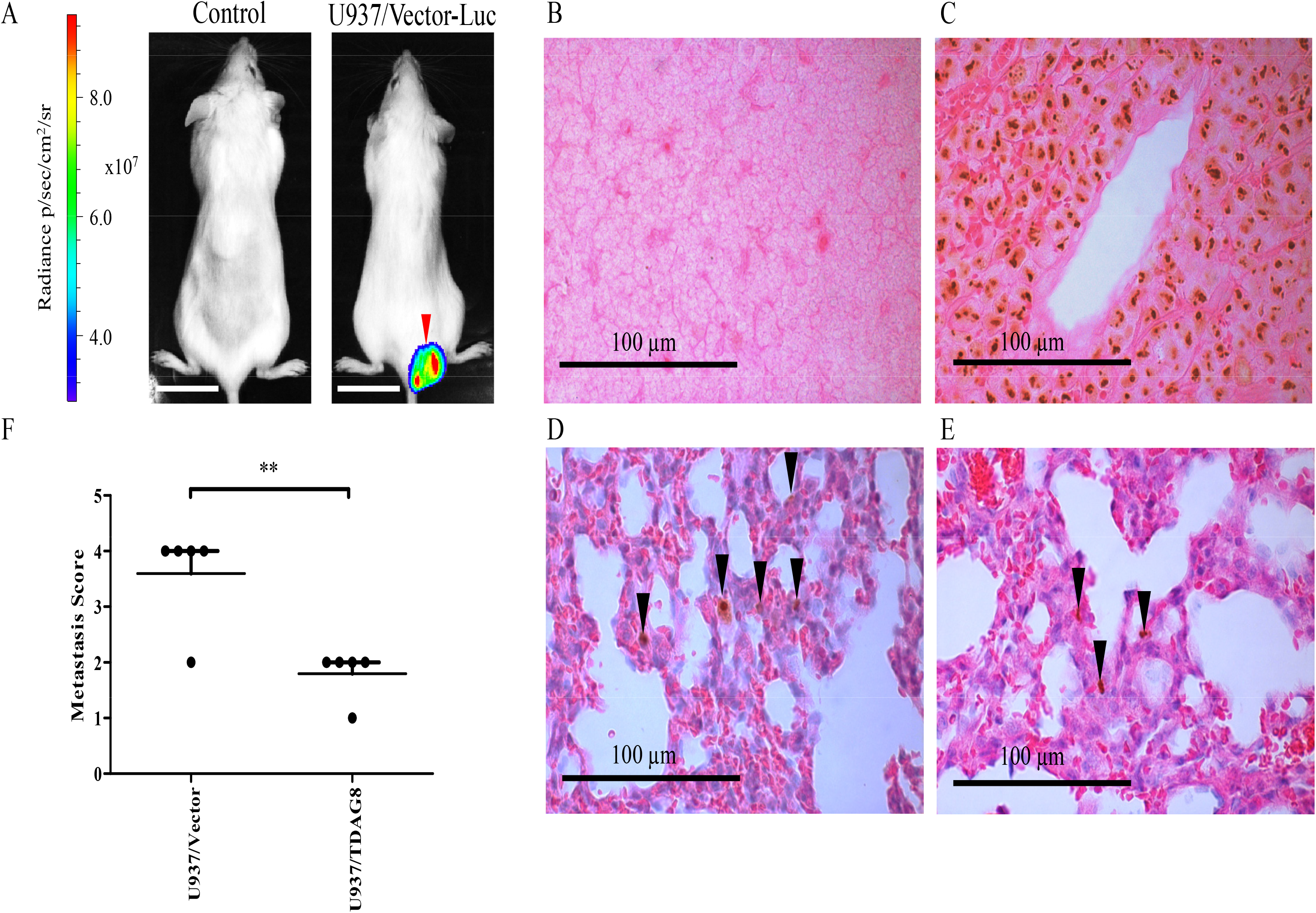
Restoration of TDAG8 gene expression reduces U937 cell metastasis in SCID mice. In vivo luminescence imaging identified larger tumors but was not sensitive enough to identify micro metastasis in SCID mice. (A) IVIS luminescence imaging. Scale bar = 2 cm. (B) Normal mouse spleen section with negative staining demonstrating the specificity of the antihuman nucleoli antibody. (C) Tumor metastasis in the lymph node was positive for human nucleoli. (D-E) Hematoxylin and eosin staining and IHC of the human nucleoli antibody was used to identify nuclear localization of the antibody signals of U937/Vector tumor cells in the lung of SCID mice. (F) Analysis of SCID mice lungs following IHC of human nucleoli reveals a significant reduction in metastasis. Mice with no metastasis detected were given a score of 0. Mice with < 2 average metastatic lesions in each 5 μm section of the lung were given a score of 1, from 2-5 average metastatic lesions were given a score of 2, from 6-10 average metastatic lesions were given a score of 3, and from 11 or more average metastatic lesions were given a score of 4. In addition, lesions that were larger than 20 cells or large enough to see with the IVIS Lumina XR unit was given a 4. For each experimental group injected with either U937/Vector-Luc or U937/TDAG8-Luc cells the individual scores were added up and averaged to give a metastasis score. The scoring system consisted of analyzing a total of 6 sections of lung for each mouse injected. Every section was separated by at least 20 μm of tissue to give representative sections of various areas of the lung. ** P < 0.01

## Discussion

The effect of extracellular acidosis on cancer progression is complex; therefore it is important to understand the cancer cell biological response to it^13^. In this study it was determined that diverse extremities of extracellular acidosis had differential effects on cellular proliferation (Figure 2C-E & G-H). It was clearly demonstrated that mild extracellular acidosis, pH 6.9, increased U937 cell proliferation while severe acidosis, pH 6.4, repressed it (Figure 2C). In addition, this report evaluated the effects of tumor acidosis on c-myc oncogene expression in vivo. In tumor sections nearest necrotic regions, known to be acidic^17^, c-myc oncogene expression was reduced significantly in live tumor cells (Figure 3D-I). However, invasive cancer cells that were invading new tissue, devoid of blood vessels and nutrients, did not follow this pattern (Figure 3D-I). Conversely, invasive tumor cells had increased expression of c-myc and Ki-67 signifying they are actively proliferating and invasive despite extracellular acidosis and nutrient deprivation (Figure 3D-I). The discovered pattern of tumor cell invasiveness in harsh extracellular conditions concurrently associating with increased cell proliferative markers is similar to an acid resistant phenotype described previously^17,18^. Importantly, understanding that extracellular acidosis reduces c-myc oncogene expression in some tumor cells while not in others provides an original understanding for the heterotypic and spatial tumor cell response to extracellular acidosis. To understand how tumor cells sense extracellular acidosis the proton sensing G-protein coupled receptor TDAG8 was investigated.

TDAG8 has been reported to have a diverse repertoire of pro- and anti-oncogenic effects that are cell type and context dependent^22–24,51–54^. The data provided in this study indicates that TDAG8 gene expression is suppressive for U937 cell malignancy. It was discovered that TDAG8 gene expression is reduced in the majority of blood cancers in comparison to normal immune cells or leukocyte-rich tissue (Table 1 & Figure 1). In addition, higher expression of the TDAG8 gene correlates with increased sensitivity toward various chemotherapeutics (Table 2). Using U937 acute myeloid leukemia cells as a model system TDAG8 gene expression was restored to a normal physiological level to test the hypothesis that TDAG8 gene expression provides a disadvantage for blood cancer cell malignancy. Restoring TDAG8 gene expression in U937 cells reduced cell proliferation in vitro and tumor growth and metastasis in vivo (Figure 2–3 & 6). The ability for TDAG8 gene expression to provide a disadvantage for U937 cell proliferation, tumor growth, and metastasis confirmed that TDAG8 gene expression provides a disadvantage for blood cancer progression. Similar tumor suppressive functions of TDAG8 were observed in other blood cancer cells such as Ramos Burkitt lymphoma and RPMI 8226 multiple myeloma cells (Supplemental Figure 3 and 5). Moreover, TDAG8’s inherent ability to reduce c-myc expression at physiological pH 7.4 and acidic pH 6.4 is central (Figure 4A-C)^23^. This is consistent with previous results demonstrating that TDAG8 exhibits constitutive activity at physiological pH and is further activated at acidic pH^41^. Moreover, the results from this report indicate that the inhibitory effects of acidosis on c-myc oncogene expression are partially due to TDAG8-mediated Gα13/RhoA signaling (Figure 4D-E). This idea aligns with recent reports that demonstrate Gα13/RhoA signaling suppresses oncogenesis and acts as a tumor suppressor in lymphoma^55,56^. Overall, it is hypothesized in this report that extracellular acidosis provides a selective pressure against cancer cells, which modulates clonal cell evolution. Moreover, TDAG8 is a proton sensor that plays an important role in this process, which has important implications for blood cancer progression as well as cancer cell clonal evolution parallel to extracellular acidosis found within the tumor microenvironment.

## Acknowledgement

We thank Dr. Owen Witte for providing the luciferase construct, Dr. Douglas Weidner for providing assistance on flow cytometry, and Dr. Zhigang Li for technical assistance at the initial stage of the project. This study was supported in part by research grants from the Vidant Cancer Research and Education Fund (to LVY), the Coach Rock Roggeman Cancer Research Fund (to LVY), and the East Carolina University Research and Creative Activity Award (to LVY).

## Authorship

Contribution: C.R.J. and L.V.Y. conceptualized and designed the study; C.R.J., E.J.S, L.D., and K.L. performed the experiments; C.R.J., E.J.S., and L.V.Y. analyzed the data; A.S. and J.T.C. performed the GEO microarray dataset analysis; C.R.J wrote the first draft of the manuscript; L.V.Y. critically revised the manuscript; all authors read and edited the manuscript; L.V.Y. acquired funding and supervised the study.

## Conflict-of-interest disclosure

The authors declare no conflict of interest.

---

**Supplemental Figure 1.**
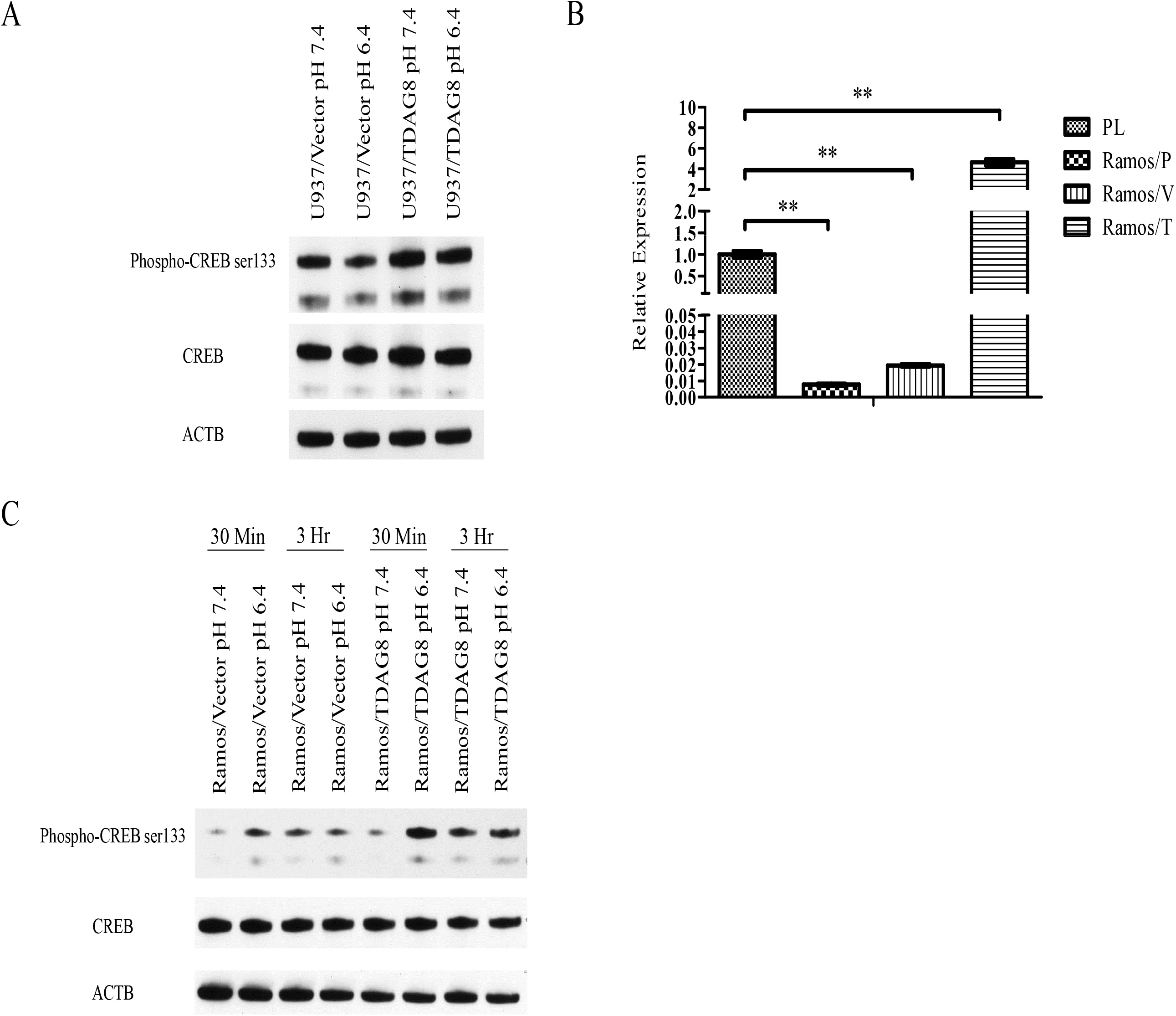
Restoration of TDAG8 gene expression results in increased phosphorylation of CREB at serine 133 in U937 and Ramos cells. (A) Restoration of TDAG8 gene expression in U937 cells results in stimulation of CREB phosphorylation at serine 133 indicating the TDAG8 activity level is increased. (B) TDAG8 gene expression is restored in Ramos cells to a level that is physiologically relevant. (C) Restoration of TDAG8 gene expression in Ramos cells results in stimulation of CREB phosphorylation at serine 133 indicating the TDAG8 activity level is increased. ** P < 0.01

**Supplemental Figure 2.**
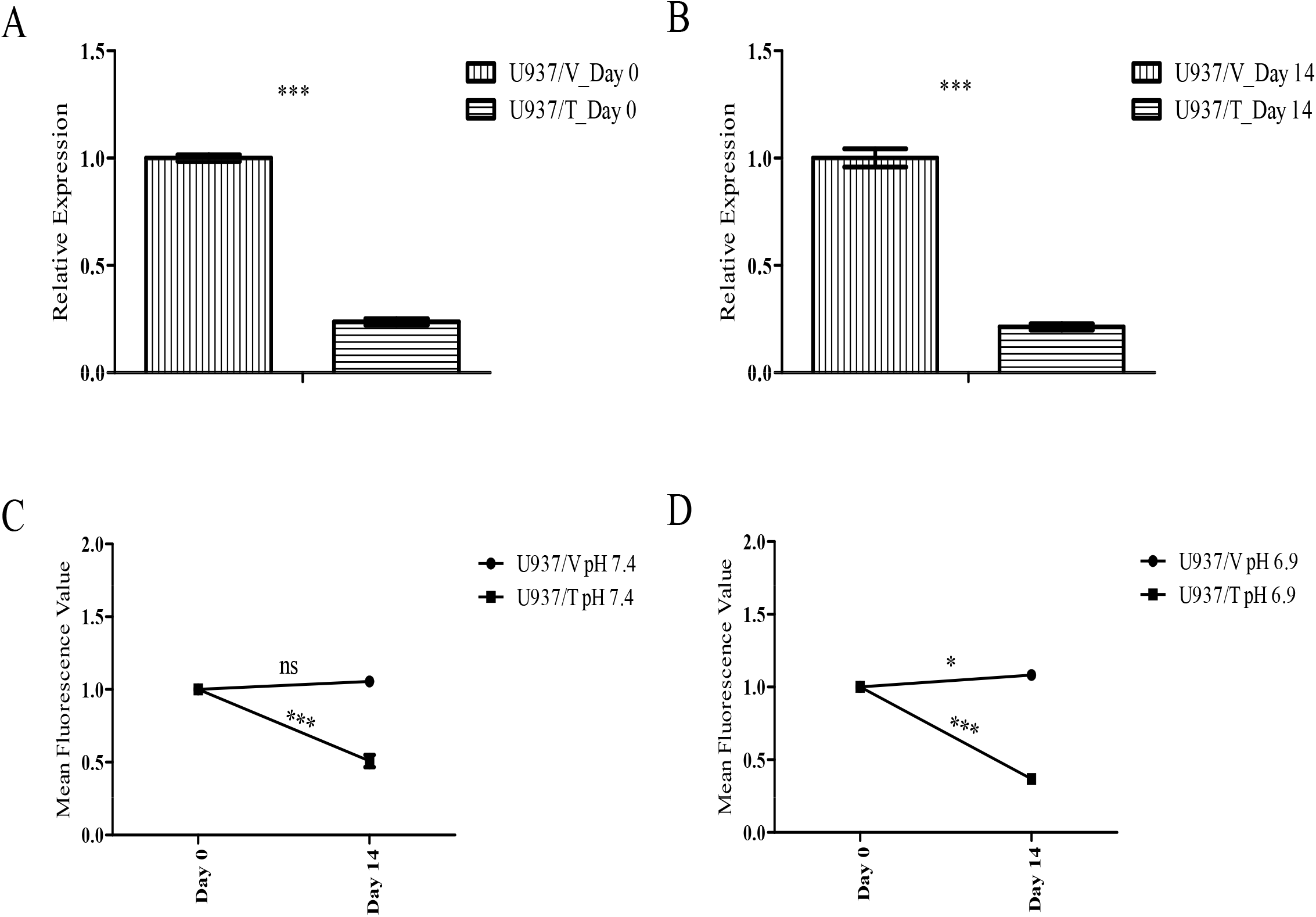
TDAG8 gene expression restoration reduces c-myc oncogene expression in U937 cells. (A-B) Restoration of TDAG8 gene expression reduces c-myc oncogene expression at the mRNA level in U937 cells. (C) Over 14 days U937/TDAG8 GFP expression is reduced at physiological pH 7.4 while U937/Vector GFP is stable. (D) Reduction of U937/TDAG8 GFP expression is further augmented by activation of TDAG8 with acidic pH 6.9 treatment while the U937/Vector GFP is stable. ns P > 0.05, * P < 0.05, *** P < 0.001

**Supplemental Figure 3.**
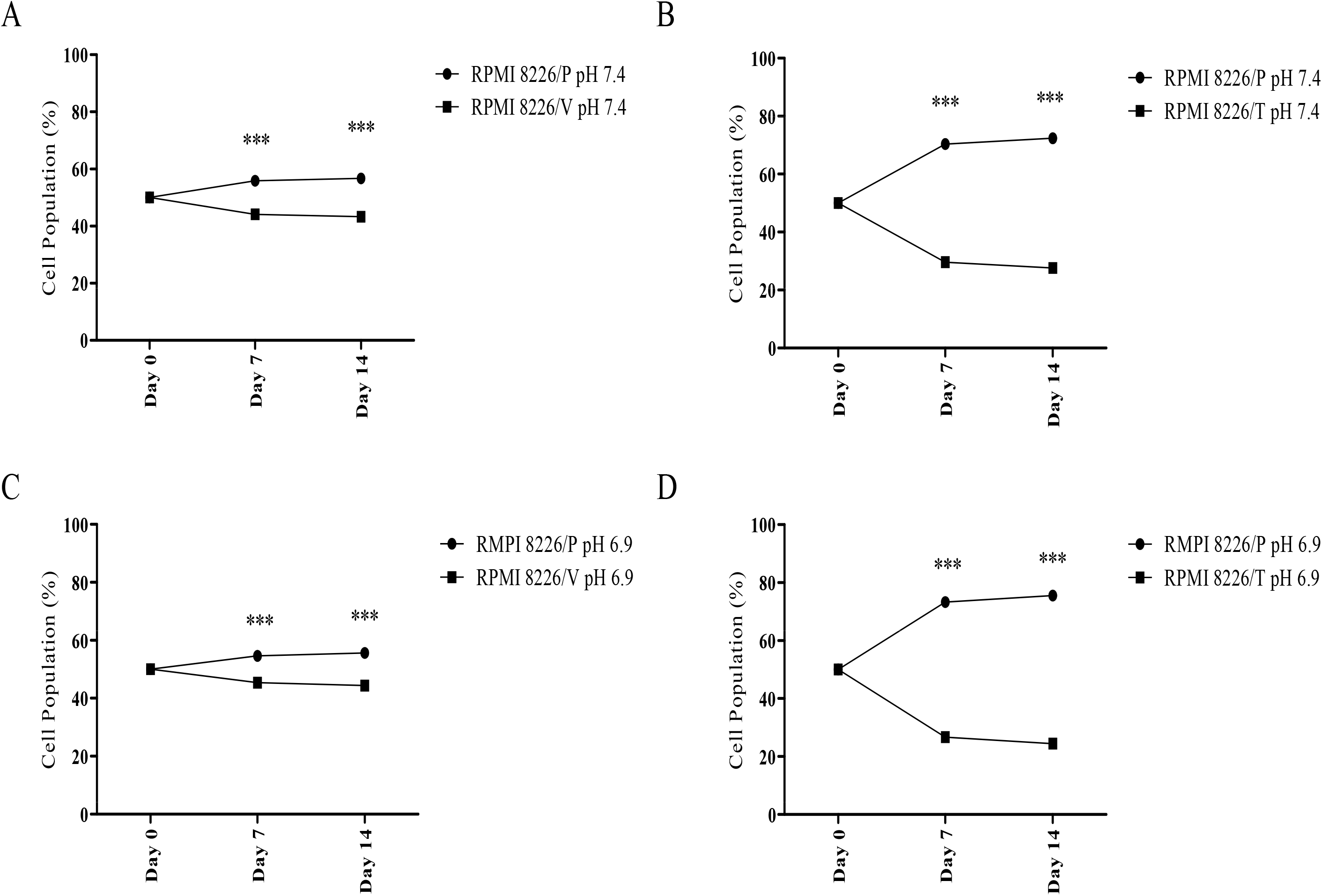
Restoration of TDAG8 gene expression in RPMI 8226 myeloma cells inhibits cell proliferation. (A) The empty vector does not substantially affect RPMI 8226 cell proliferation at physiological pH 7.4 in comparison to the RPMI 8226 parental cells. (B) Restoration of TDAG8 gene expression significantly reduces RPMI 8226 cell proliferation at physiological pH 7.4 in comparison to the RPMI 8226 parental cells. (C) The empty vector does not substantially affect RPMI 8226 cell proliferation at acidic pH 6.9 in comparison to the RPMI 8226 parental cells. (D) Restoration of TDAG8 gene expression significantly reduces RPMI 8226 cell growth at acidic pH 6.9 in comparison to the RPMI 8226 parental cells. *** P < 0.001

**Supplemental Figure 4.**
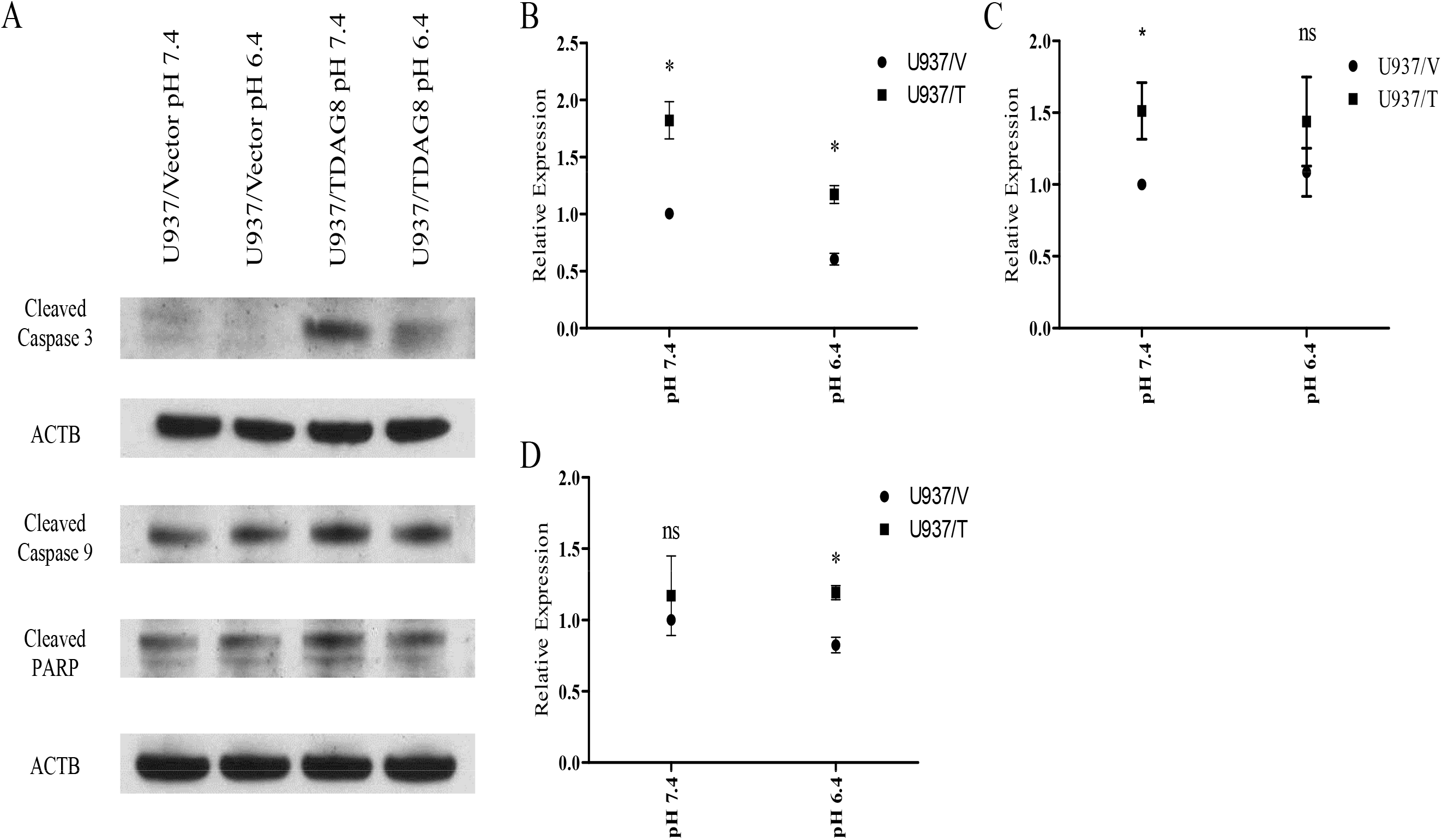
Restoration of TDAG8 gene expression increases apoptosis signaling. (A-B) Restoration of TDAG8 gene expression stimulates cleaved caspase 3 in U937 cells at physiological pH 7.4 and acidic pH 6.4. (A & C) Restoration of TDAG8 gene expression increases cleaved caspase 9 in U937 cells at physiological pH 7.4. (A & D) Restoration of TDAG8 gene expression increases cleaved PARP in U937 cells at acidic pH 6.4. ns P > 0.05, * P < 0.05

**Supplemental Figure 5.**
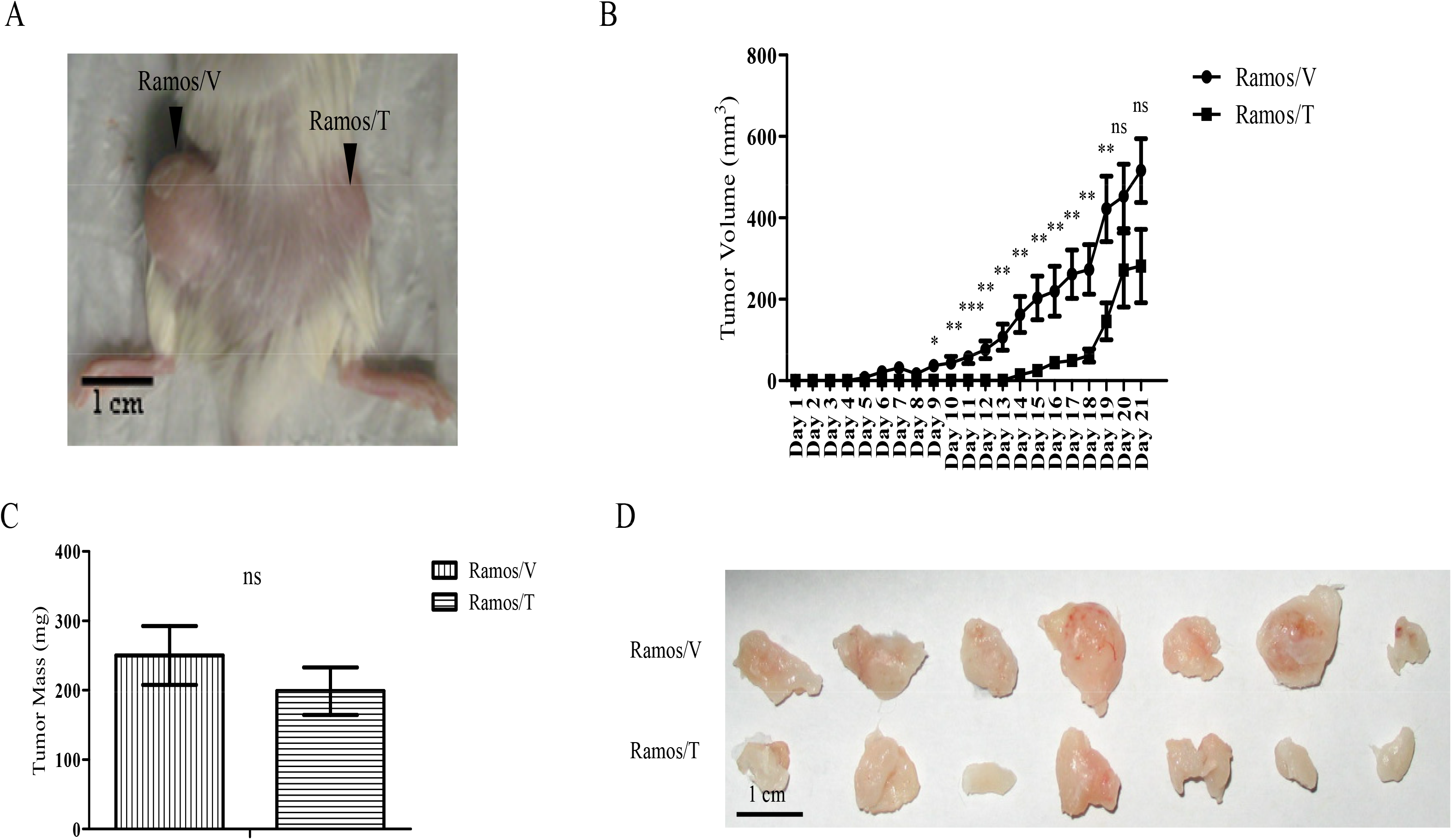
Restoration of TDAG8 gene expression in Ramos lymphoma cells reduces primary tumor growth in SCID mice. (A) Representative image of a mouse subcutaneously injected with Ramos/Vector (left flank) and Ramos/TDAG8 (right flank) cells. (B) Restoration of TDAG8 in Ramos cells significantly delays primary tumor growth in SCID mice starting day 9 after injection. (C) Restoring TDAG8 gene expression in Ramos cells moderately reduces overall tumor mass after necropsy on day 21. (D) Representative image of Ramos/Vector and Ramos/TDAG8 tumors excised from SCID mice. ns P > 0.05, * P < 0.05, ** P < 0.01, *** P < 0.001

**Supplemental Figure 6.**
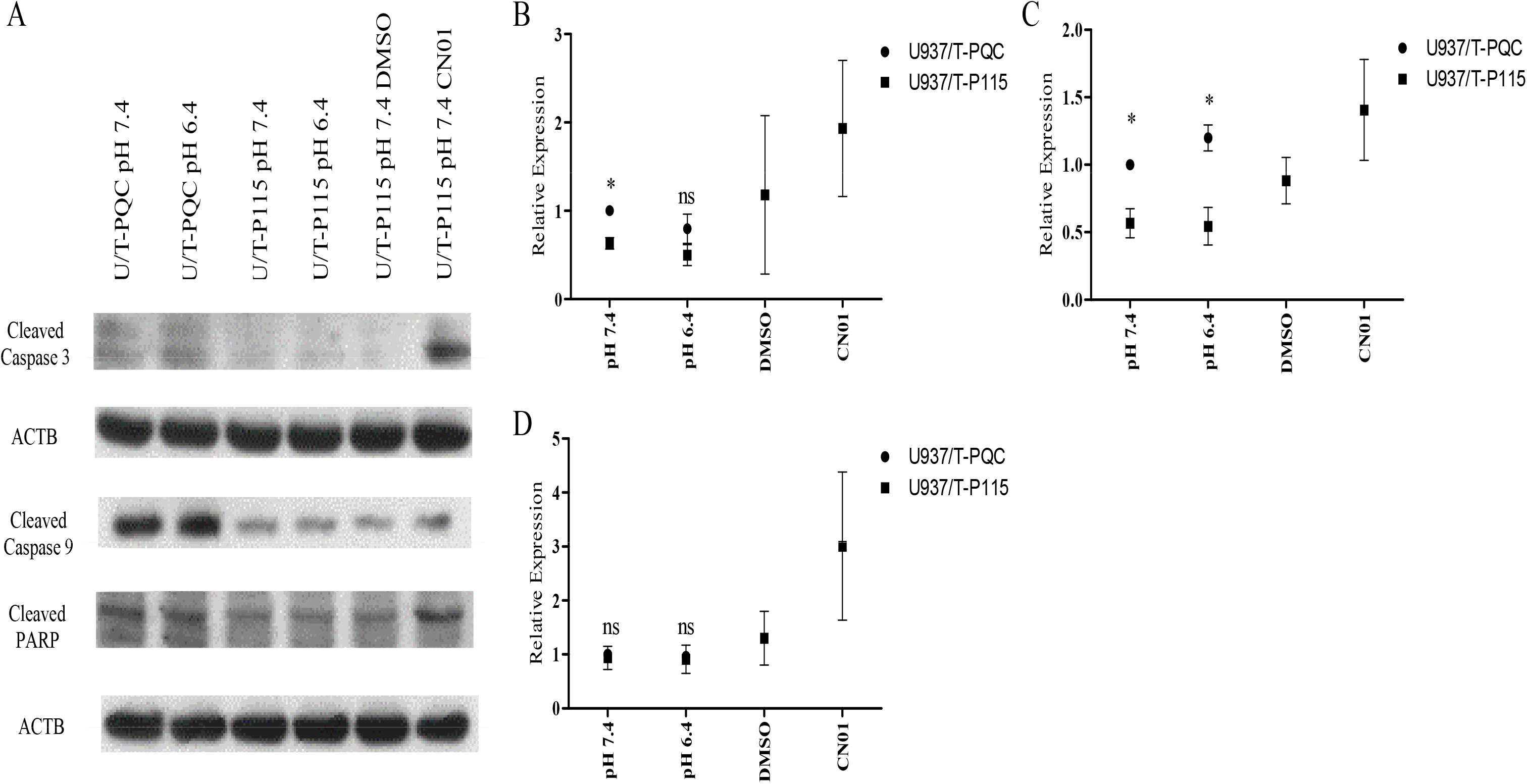
TDAG8 stimulates apoptotic signaling through Gα13/RhoA signaling. (A-B) Inhibition of Gα13 signaling in U937/TDAG8 cells reduces cleaved caspase 3 and activation of RhoA in U937/TDAG8 cells stimulates cleaved caspase 3. (A & C) Inhibition of Gα13 signaling in U937/TDAG8 cells reduces cleaved caspase 9 and activation of RhoA in U937/TDAG8 cells stimulates cleaved caspase 9. (A & D) Inhibition of Gα13 signaling in U937/TDAG8 cells reduces cleaved PARP and activation of RhoA in U937/TDAG8 cells stimulates cleaved PARP. ns P > 0.05, * P < 0.05

**Supplemental Figure 7.**
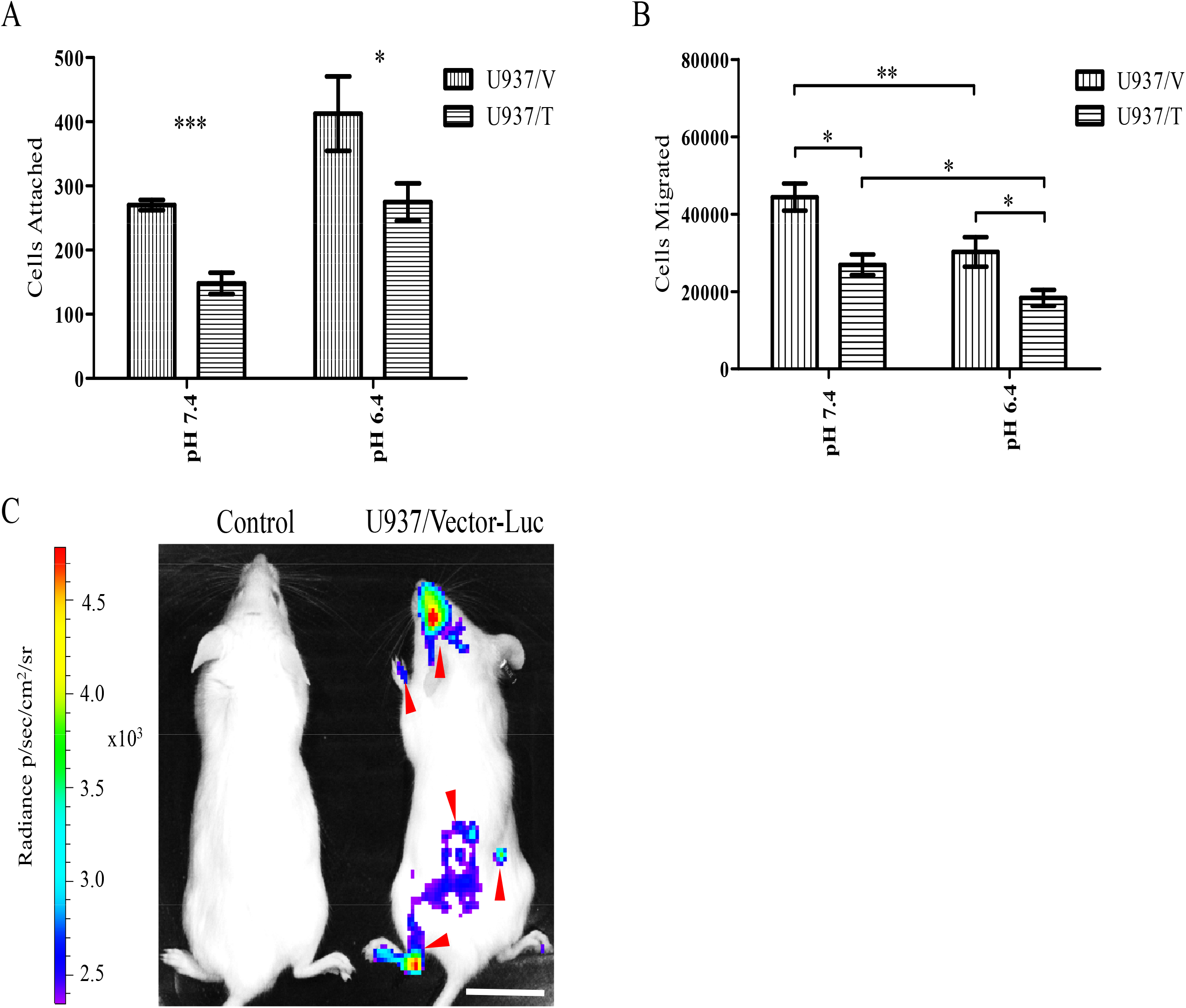
Restoration of TDAG8 gene expression in U937 cells reduces attachment to a HUVEC monolayer and reduces migration toward a chemoattractant. (A) Restoration of TDAG8 gene expression reduces overall U937 cell attachment to a HUVEC monolayer while extracellular acidosis increases cell attachment. (B) Restoration of TDAG8 gene expression significantly reduces U937 cell migration toward a chemoattractant (SDF-1α) while extracellular acidosis reduces overall U937 cell migration. (C) In vivo imaging of a SCID mouse injected with U937/Vector-Luc cells that presented with hind limb paralysis and metastasis throughout the body. * P < 0.05, ** P < 0.01, *** P < 0.001

